# Orientation-locked DNA origami for stable trapping of small proteins in the NEOtrap

**DOI:** 10.1101/2022.09.09.507286

**Authors:** Chenyu Wen, Eva Bertosin, Xin Shi, Cees Dekker, Sonja Schmid

## Abstract

Nanopores are versatile single-molecule sensors that offer a simple label-free readout with great sensitivity. We recently introduced the Nanopore Electro-Osmotic trap (NEOtrap) which can trap and sense single unmodified proteins for long times. The trapping is achieved by the electro-osmotic flow (EOF) generated from a DNA-origami sphere docked onto the pore, but thermal fluctuations of the origami limited the trapping of small proteins. Here, we use site-specific cholesterol functionalization of the origami sphere to firmly link it to the lipid-coated nanopore. We can lock the origami in either a vertical or horizontal orientation which strongly modulates the EOF. The optimized EOF greatly enhances the trapping capacity, yielding reduced noise, reduced measurement heterogeneity, an increased capture rate, and 100-fold extended observation times. We demonstrate the trapping of a variety of single proteins, including small ones down to a molecular mass of 14 kDa. The cholesterol functionalization significantly expands the application range of the NEOtrap technology.

---

Few techniques have the ability to sense single biomolecules in a label-free manner, and even fewer can do so in solution and at room temperature. The recently developed Nanopore Electro-Osmotic trap (NEOtrap) is such a label-free single-molecule technique that can trap and study proteins one by one.^1,2^ As shown in Figure 1a, the NEOtrap consists of a DNA-origami sphere that is docked onto a passivated solid-state nanopore when a positive bias voltage is applied (to the trans side). Once docked, the highly negatively charged DNA-origami sphere generates an electro-osmotic flow (EOF) by which a target protein can be trapped (Figure 1b). Various protein-specific features, such as protein size and distinct conformations, can be monitored by recording the through-pore ion current. This electrical readout provides the NEOtrap with a broad temporal bandwidth compared to other single-molecule techniques:^3^ big proteins, such as ClpP (340 kDa), in suitable conditions (proper pore size and bias voltage) can be trapped and observed for up to hours with a time resolution of microseconds.^1^ However, it appeared challenging to trap small proteins for a long time in the NEOtrap. As shown in Figure 1c, avidin (54 kDa; ~6 nm in diameter^4^) exhibits a typical trapping time of only milliseconds. We hypothesized that the short trapping time is likely limited by thermal positional fluctuations of the DNA-origami sphere allowing the through-pore escape of the protein. This led us to a new NEOtrap design which we report in the current Letter.

**Figure 1.**
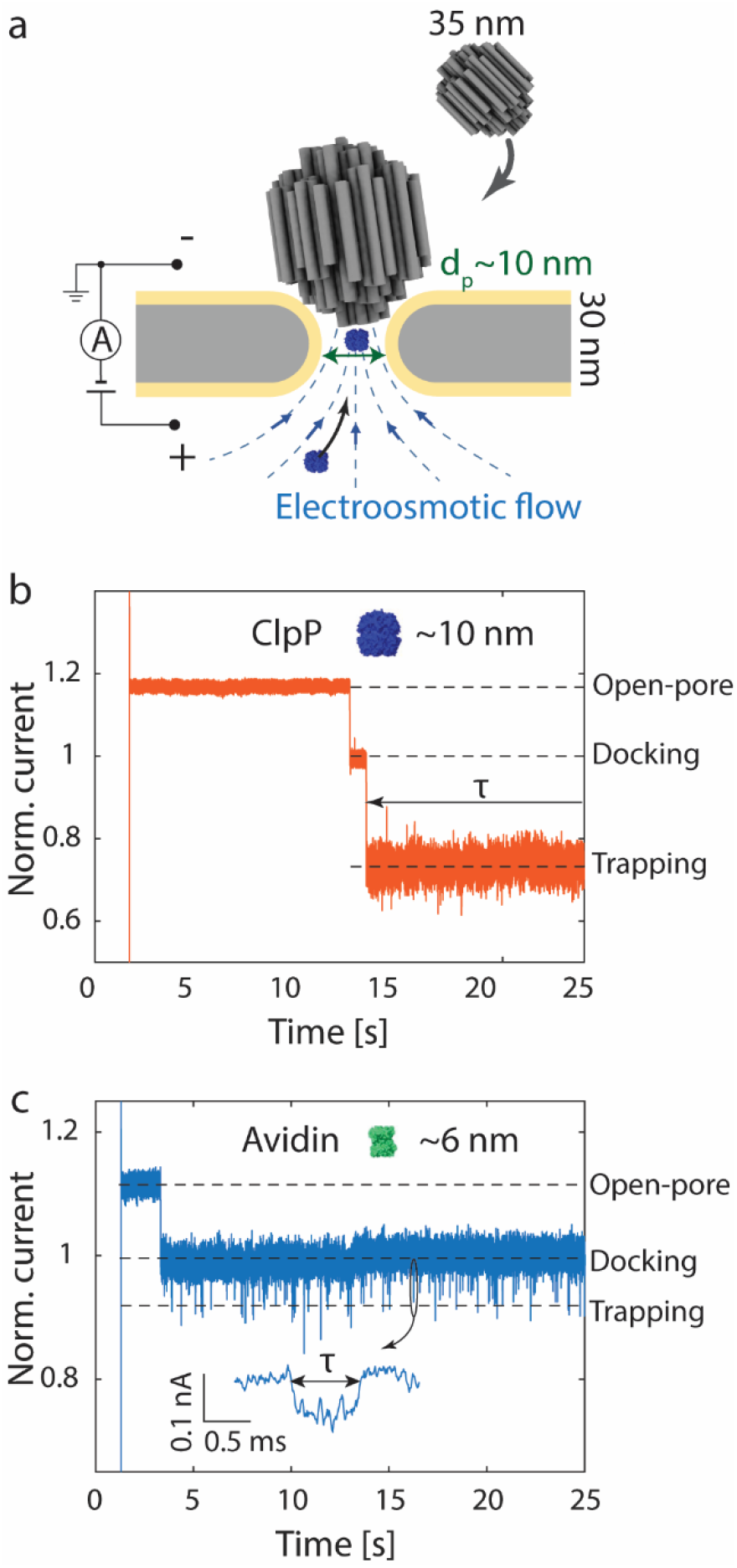
Protein trapping with the NEOtrap. (a) Schematic of the NEOtrap. A solid-state nanopore is coated with a lipid bilayer for passivation to prevent the non-specific adsorption of biomolecules. Under a positive bias voltage at the trans side, a DNA-origami sphere at cis is attracted and docked onto the pore. This generates an EOF through the porous structure of the origami sphere. this EOF attracts a target protein from the trans side and traps it in the pore. (b) Typical trapping current trace of a ClpP protein in a 10 nm nanopore (after lipid bilayer coating). Upon applying a 120mV voltage, the open pore current is measured, followed by a current drop caused by the docking of the DNA-origami sphere, and a second current drop indicating the trapping of the ClpP protein. (c) Same as (b) but for avidin in a ~10.5 nm nanopore. Inset shows a zoom of a typical trapping event. Protein Data Bank (PDB) codes ClpP, 1yg6; avidin, 3mm0.

Here, we present the ‘NEOtrap 2.0’ with significantly enhanced trapping and sensing performance down to small proteins, which is achieved by two advances: first, we block the through-pore escape pathway by firmly attaching the DNA-origami sphere to the nanopore. Second, guided by numerical simulations, we conceived a new way to tune the magnitude of the EOF *in-situ*, namely by controlling the orientation of the docked origami sphere. We realized this orientation locking experimentally and show a strong orientation-dependence of the EOF in protein trapping experiments. Finally, we demonstrate the enhanced sensing performance of the ‘vertically’ orientation-locked NEOtrap, based on a variety of proteins in a size range down to 13.7 kDa. Remarkably, we find a 100-fold increase of the trapping times, more homogenous trapping, and significantly reduced noise compared to the previous design.

To improve the trapping capacity of the NEOtrap, we locked the DNA-origami sphere onto the lipid bilayer-coated nanopore by attaching cholesterol molecules to its surface. We coupled twelve cholesterols to the origami, one at each of the twelve corners of the icosahedral origami structure (see Methods). Cholesterol molecules are known to insert into lipid bilayers owing to their strong hydrophobic interaction with amphiphilic molecules. Here, they thus act as anchors that firmly attach the sphere onto the nanopore (see Figure 2a). This can be observed in our experiments by comparing the current recordings for cholesterol-functionalized origami versus traces for bare spheres: without functionalization (Figure 2b), application of a negative bias ejected the negatively charged origami and the open-pore current was recovered when flipping back to positive voltage, shortly thereafter followed by a new origami docking event. By contrast, the cholesterol-functionalized DNA-origami sphere (Figure. 2c) was quasi-permanently attached to the nanopore upon its first docking, and it stayed firmly docked during repeated voltage inversions, leading to the observed constant current levels in Figure 2c.

**Figure 2.**
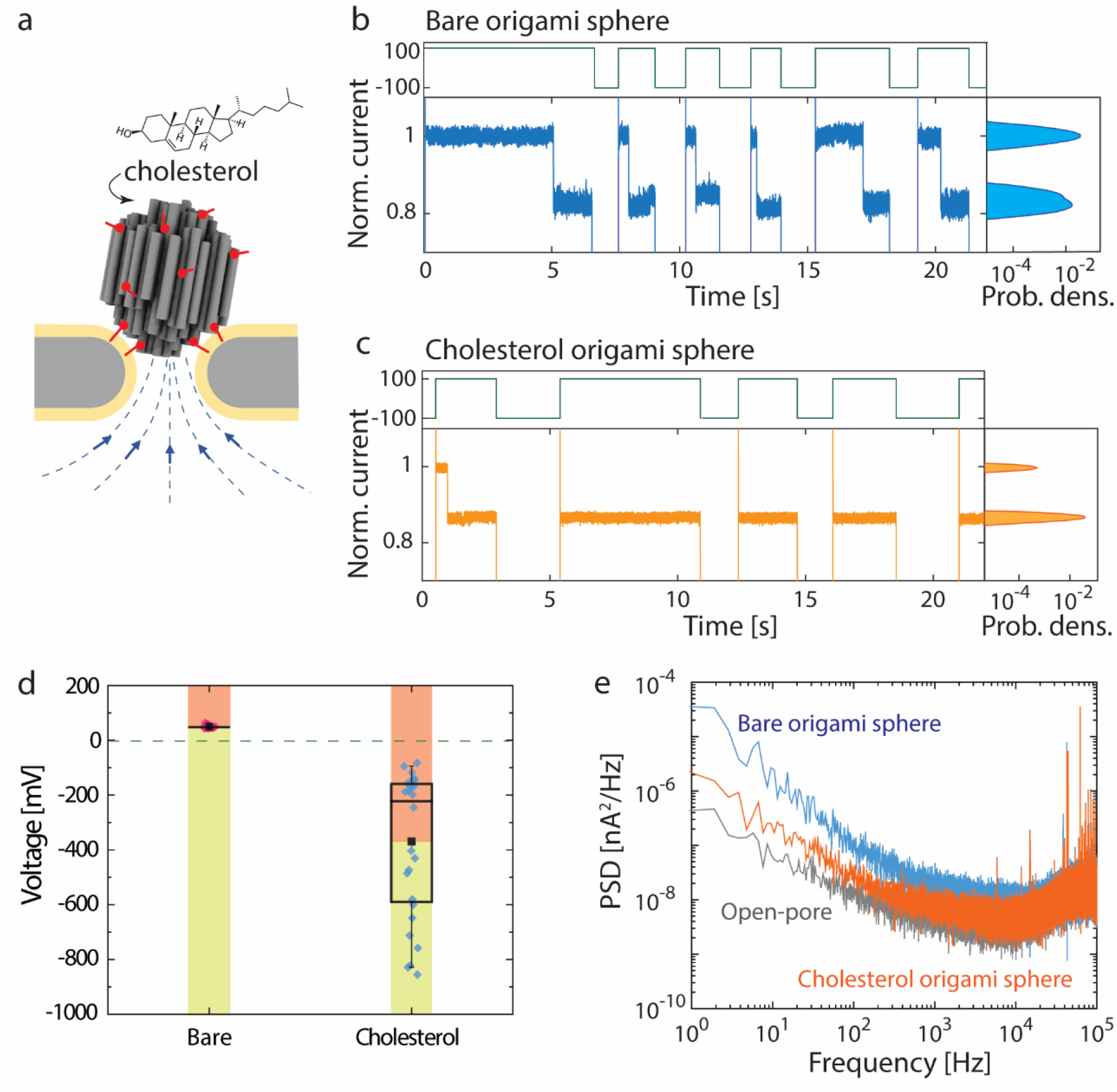
Docking of cholesterol-functionalized DNA-origami spheres. (a) Schematic showing the anchoring of the DNA-origami sphere on the lipid bilayer by cholesterol molecules on the origami sphere. (b, c) Current traces showing the docking of the bare spheres (b) and the cholesterol-functionalized sphere (c), as demonstrated by changing the voltage polarity multiple times. The Histograms of the current traces are shown on the right. The alternation of the corresponding added voltage between −100 mV and 100 mV is shown at the top of the current traces. (d) Box chart comparing the distribution of voltages for release a docked bare origami sphere and cholesterol-functionalized origami sphere. In one box, the dot shows the average value, the line in the box indicates the median, upper and lower boundaries represent the 25% and 75% data, and the error bar marks the 5%-95% range. (e) Power spectrum density of the baseline noise under different conditions.

We then tested at which voltage the DNA-origami sphere detached from the pore, using voltage ramps from positive to negative bias (See SI Note 1, Figure S1-S3). As shown in Figure 2d, without cholesterol functionalization, the DNA-origami sphere detached already at small positive voltages (of about +50 mV), where thermal fluctuations were sufficient to overcome the electrophoretic docking potential. By contrast, the cholesterol-functionalized sphere stayed attached even up to considerable negative voltages, before detachment at a voltage between −200 to −800 mV. At these high applied voltages, the lipid bilayer coating got destabilized, causing increased noise. In some cases, the lipid bilayer was even totally peeled off (See SI Figure S4), which suggests that the cholesterols on the origami sphere pulled out lipids from the bilayer during rupture.

Gratifyingly, the cholesterol-functionalized spheres showed significantly reduced current noise, with typical values of *I_RMS_* = 8pA, which is close to the open-pore baseline of the same lipid-coated pore (*I_RMS_* = 7pA), and ~40% less than that for non-functionalized spheres (*I_RMS_* = 13pA, all measured at 100mV and 10 kHz bandwidth). As shown in the power spectrum in Figure 2e, this noise reduction originates predominantly from a low-frequency 1/f-type noise (<1 kHz) which is reduced by ca. an order of magnitude upon cholesterol functionalization. This can be attributed to reduced thermal fluctuations, as 1/f low-frequency noise has been ascribed to mechanical instabilities.^5^ Altogether, the cholesterol functionalization strongly reduces the excess noise – almost to the open-pore level.

We found that the orientation of the DNA-origami sphere on top of the nanopore is of great interest. Notably, the ‘sphere’ is only pseudo-isotropic, as it is built of many parallel DNA helices (cf. Figure 2a). Accordingly, it can dock onto the pore in various orientations. To estimate the effect of such different origami orientations on the EOF, we first performed finite-element simulations using the COMSOL Multiphysics software. We simulated our experimental conditions using an axis-symmetrical two-dimensional approximation, with an origami sphere mimicked using ‘DNA rods’, void nanochannels, and corresponding parameters. As illustrated in Figure 3a, we considered the two extreme cases of a vertically or horizontally oriented sphere, where the DNA helices are parallel or perpendicular to the pore axis, respectively. The electric field, ion transport, and water movement, were fully coupled, as described by the Poisson, Nernst-Planck, and Navier-Stokes equations, respectively^6^ (see Methods for details). These simulations yielded the EOF fields and the water and ion flows, and Figure 3a shows the substantially different EOF distribution for both sphere orientations. Clearly, vertical sphere docking causes a much stronger EOF through the nanopore, compared to the horizontal configuration, leading to a water velocity for the vertical docking that is significantly higher than that for the horizontal docking (factor 2.5 times in our approximation, see Figure 3b). This can be intuitively understood as a result of a less obstructed EOF in the vertical case. Such an increased EOF for the vertical docking should lead to experimentally measurable effects in protein trapping experiments, e.g., an increased capture rate. Therefore, to test these results experimentally, we went on to realize specific vertical and horizontal docking orientations.

**Figure 3.**
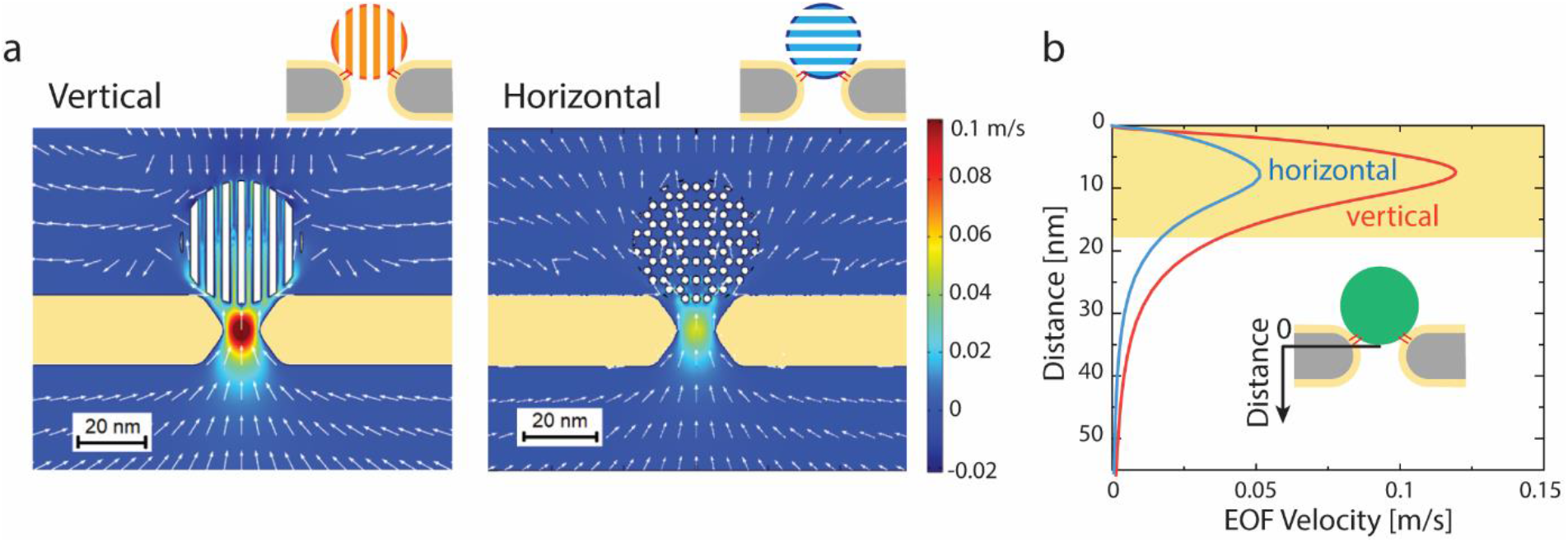
Simulation results of the NEOtrap. (a) Distribution of EOF generated by the vertically and horizontally docked origami spheres. Color represents the vertical component of the flow velocity with the positive direction pointing upwards, and the arrows indicate the flow direction. (b) Velocity of the water flow along the central axis as marked in the inset. The distance counts from the bottom of the origami sphere pointing downwards.

We realized orientation-locked docking of the DNA-origami sphere on the lipid-coated nanopore (Figure 4a), by attaching the cholesterol molecules at specific locations of the sphere, instead of the uniform distribution discussed above. In the ‘vertical design’, cholesterol molecules were attached only at one end of the DNA double-helices, whereas the ‘horizontal design’ featured cholesterols at one side plane of the DNA origami spheres. The two orientations of docking showed distinct current blockades: the vertical orientation generated a deeper relative blockage (17% of the open-pore current) compared to the horizontal counterpart (14.5% of the open-pore current; both measured on the same pore), see Figure 4b and SI, Figure S5. We attribute this to the non-isotropic sphere geometry which allows the vertically oriented sphere to reach deeper into the pore, and thus block more through-pore current, than the horizontally oriented sphere. In the docking experiments, we typically observed that voltage inversions first led to sphere release and baseline recovery for a few times, until a final permanent (i.e. voltage-inversion-resistant) origami sphere arrangement was realized with the designed orientation on the nanopore, see for example Figure 4b. The vertical design was observed to need fewer attempts than the horizontal design to achieve the permanent docking arrangement.

**Figure 4.**
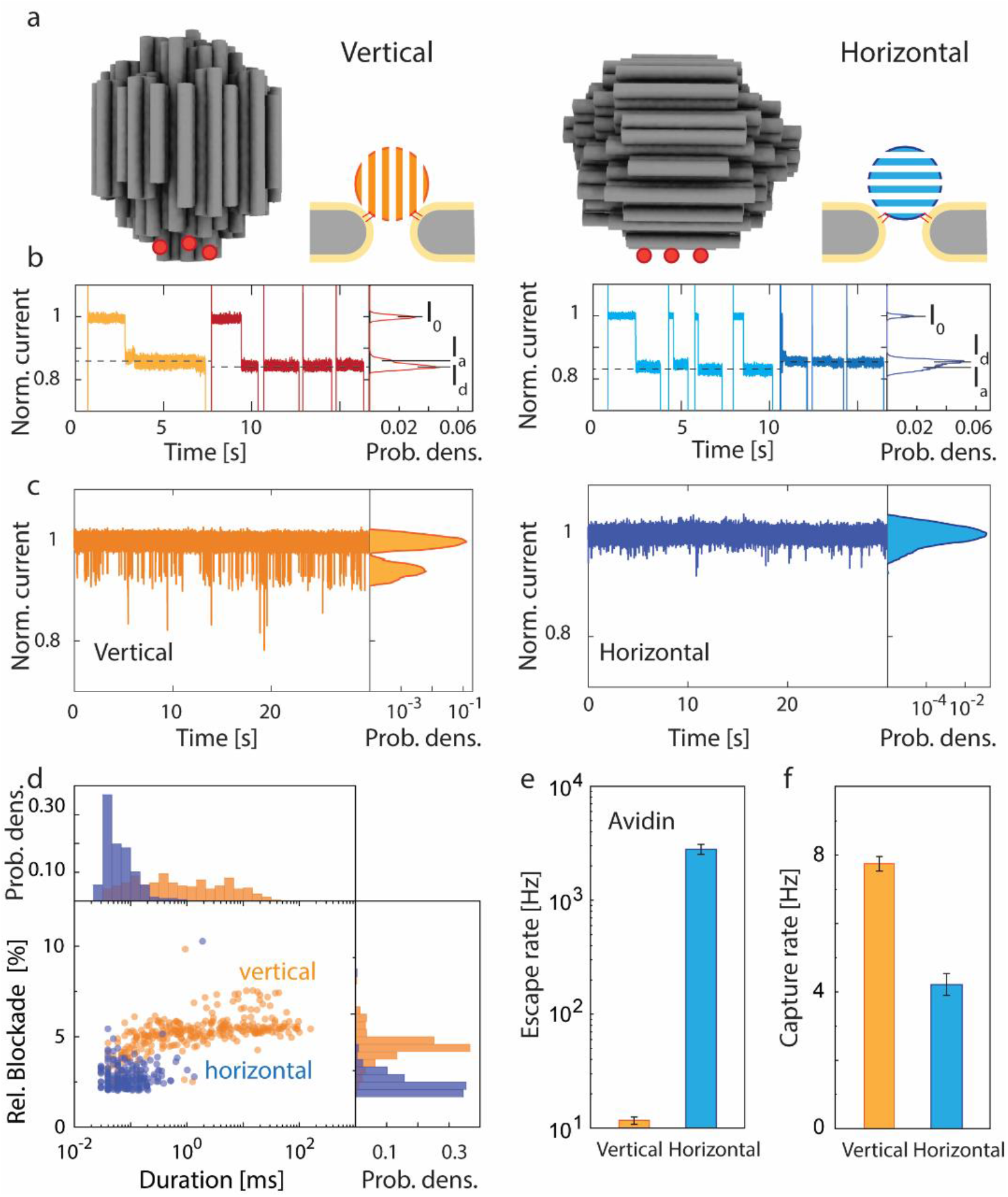
Influences of orientation of the DNA-origami spheres on protein trapping. (a) Schematic showing the position of the functionalized cholesterol on the DNA-origami sphere for vertical (left) and horizontal (right) configurations. Red dots indicate the positions of the added cholesterol molecules. (b) Current traces of docking of DNA-origami spheres with a vertical (left) or horizontal (right) configuration. The traces show that the spheres need several attempts (light orange/blue colors) before landing on the nanopore with the final designed orientation that is locked by the cholesterol anchors (dark red/blue colors). Histograms of the corresponding current traces are shown on the right-hand side. Currents are normalized by the open pore current (*I*_0_). For the vertical configuration, the optimal docking that locked by the cholesterol anchors generated a deeper final blockage level (*I*_d_), compared to the level of attempt dockings (*I*_a_), while *I*_d_ is higher than *I*_a_ for the horizontal configuration. (c) Current traces for trapping of avidin by vertically (left) and horizontally (right) docked origami spheres at 100 mV bias. Histograms of the corresponding current traces are shown on the right-hand side. (d) Scatter plots compare the trapping time and relative blockage amplitude of the trapping events for avidin. Comparison of escape rate (e) and capture rate (f) of avidin by vertical and horizontal origami spheres configurations at 100 mV bias. The error bars show the standard deviation of extracted parameters from fitting by using bootstrap sampling.

Next, we tested the protein trapping properties of the vertically and horizontally orientation-locked spheres. Indeed, the two docking orientations led to significantly different trapping observations, as shown in the current traces for avidin trapping in Figure 4c. In line with our simulation results, the vertical docking showed a higher capture rate (7.7 molecules/s) and a reduced escape rate (11 molecules/s), leading to more frequent and longer trapping events, as compared to those of the horizontal docking. From the scatter plot (Figure 4d), it is clear that the vertical configuration can trap avidin with a well-defined deeper blockade and for a much longer time up to ~100 ms, as compared to the ~0.1 ms in the horizontal case. Notably, the same trends are found for other proteins, such as ovalbumin (see SI, Figure S6). As quantified in Figure 4e, we consistently found that the vertical configuration showed a significantly faster capture rate (by a factor of 2) and a slower escape rate (factor of 270), leading to favorable longer trapping and sensing times. The fact that the vertical configuration gave similar trapping times as those found for spheres without specific orientation locking (i.e., with the uniformly cholesterol-functionalized DNA-origami spheres: 50 ms for avidin at 100 mV) indicates that the vertical orientation is the naturally preferred docking orientation. We suggest that this tendency to orient the origami sphere vertically onto the nanopore is caused by a geometric alignment of the origami’s DNA helices with the electric field (consistent with an anisotropic ion mobility found in DNA-origami structures^7^) while the sphere approaches the nanopore electrophoretically.

The experimental protein trapping results strongly support the notion of an orientation dependence of the EOF, as proposed by the numerical simulations. The increased capture rate and reduced escape rate indicate a larger hydrodynamic trapping potential due to more EOF if the DNA-origami is vertically oriented rather than horizontally. This can be understood by considering the microscopic structure of the DNA-origami sphere where DNA helices are arranged in a honeycomb lattice that define intermediate nanochannels of 1-2 nm in diameter. In the vertical configuration, these nanochannels are aligned with the main direction of the EOF, which is caused by the positive bias voltage acting on the counter cations that screen the DNA’s negative charge. The electrical field drives these accumulated cations upwards along the nanochannels, which drags water molecules along to form the EOF. However, in the horizontal configuration, the nanochannels are aligned perpendicularly and thus impede the vertical ion and water flow, adding additional friction. In addition, in the vertical configuration, the DNA-origami structure reached deeper into the pore (as shown by the deeper current blockade in Fig. 4b), and thus the electro-osmotically active DNA material got exposed to stronger electric fields causing more hydrodynamic flow. The viscous shear force acting on a trapped protein in this vertical case can be estimated to be on the order of magnitude of few pN (SI Note 2). Altogether, the vertical orientation-locking presents enhanced EOF and improved trapping properties for the NEOtrap.

Lastly, we directly compared the new vertically orientation-locked design introduced here with the original NEOtrap design. Exploring a set of six proteins of varied sizes, we found that the vertically docked spheres very significantly improved the trapping performance of the NEOtrap by prolonging the trapping time, as shown in Figure 5a. For example, trapping events of avidin, ovalbumin, and carbonic anhydrase showed a 10 ms – 100 ms trapping time by using the vertically orientation-locked origami spheres, while it was shorter than 1 ms using the previous non-functionalized origami design. Consequently, the vertically locked spheres provide more observation time for dynamic processes that occur in many protein systems.^8^ Furthermore, owing to the reduced thermal noise by the cholesterol-induced linkage to the pore, the events show an improved signal-to-noise ratio, and we also found a better reproducibility from experiment to experiment due to the better-defined orientation-locked configuration.

**Figure 5.**
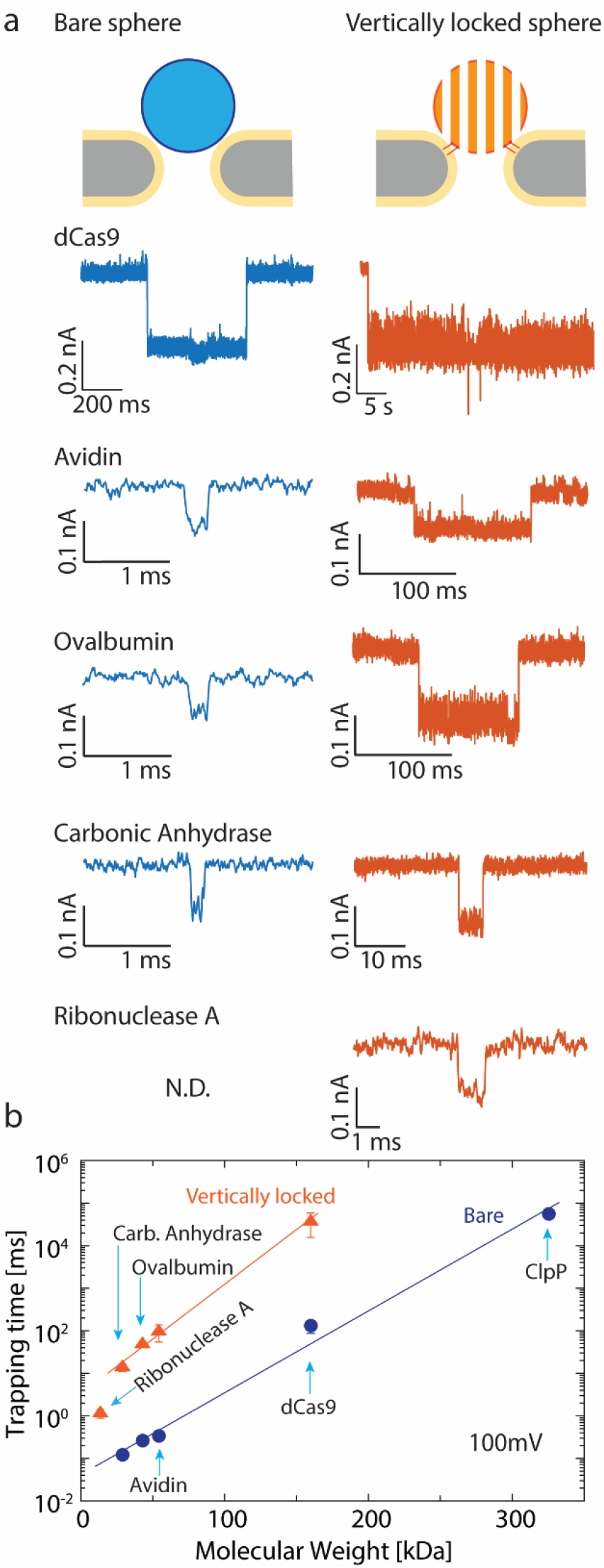
Trapping of protein by using vertical functionalized DNA-origami sphere. (a) Typical examples of trapping events of the five different proteins by using bare origami sphere and vertical functionalized origami sphere. Note the vastly expanded time scale on the right panels, which indicates the strongly enhanced stability of the trapping. The associated current traces are found in the SI (Figure S9). (b) Trapping time for different proteins versus their molecular mass, comparing bare and cholesterol-functionalized origami spheres at 100 mV bias. The error bars show the standard deviation of extracted parameters from fitting by using bootstrap sampling.

To quantify the achieved longer trapping times, we compiled the trapping time histograms, and fitted them with an exponential to extract the characteristic trapping time, i.e., the inverse escape rate. For avidin, the cholesterol-functionalization resulted in a 98 ± 8 fold increase of the trapping time for various bias voltages (See SI Figure S7). We examined the trapping of six different proteins with a mass ranging from 13.7 to 340 kDa: ribonuclease A (13.7 kDa), carbonic anhydrase (29 kDa), ovalbumin (43 kDa), avidin (54.3 kDa), dCas9 (160 kDa), and ClpP (340 kDa). An exponential dependence of the trapping time on molecular weight was obtained, in agreement with previous results.^1^ We consistently observed that the cholesterol origami spheres offered 100-fold longer observation times of unmodified proteins, as summarized in Figure 5b. We attribute this to the cholesterol-induced pore linkage which blocks the through-pore escape of protein. The observed systematic increase of the trapping time for all proteins by about 2 orders of magnitude converts to an increase of the trapping potential/energy barrier of ~5 k_B_T (see SI Note 3). The smallest protein that we examined for trapping with the vertically locked sphere was 13.7 kDa (Ribonuclease A), while, notably, such small proteins could not be trapped without cholesterol functionalization (see SI, Figure S8).

In this Letter, we presented a NEOtrap 2.0 system with a strongly enhanced trapping capacity, including reduced noise, reduced measurement heterogeneity, increased capture rate, 100-fold extended observation time, and last but not least an increased detectable range of protein masses down to 14 kDa. This was achieved by site-specific cholesterol functionalization, which locks the DNA-origami sphere in a defined vertical orientation onto the nanopore. Damping the thermal fluctuations of the sphere thus reduced low-frequency noise, and the tight linkage to the nanopore minimized through-pore escape routes. As shown by simulations and experimental data, the vertically locked sphere showed an enhanced EOF. Stabilization of docking and control of EOF were realized by the cholesterol anchors, which extend the applicable range of the NEOtrap to smaller proteins. The added control obtained with this new NEOtrap significantly improved the reproducibility and uniformity of the results, paving the way for more complex biological studies in the future.

## Methods

### Nanopore fabrication and ionic current measurement

The nanopores were drilled by a transmission electron microscope (Titan aberration-corrected TEM, Thermo Fisher Scientific, USA) in freestanding 20-nm-thick SiN_x_ membranes deposited on glass chips as described previously.^9^ Glass chips were purchased from Goeppert LLC. (USA). Nanopore devices were rinsed with deionized water (DIW, Milli-Q^®^, Merck KGaA, Germany), acetone, ethanol, isopropanol, and DIW, in sequence as mentioned. Afterwards, they were further cleaned by oxygen plasma (SPI Supplies^®^ Plasma Prep III^™^, USA) before mounted in a custom-made polyether ether ketone (PEEK) flow cell with an electrolyte reservoir at each side of the nanopore and corresponding fluidic tubing. The entire setup was placed in a Faraday cage for screening the electromagnetic interference for the electrical measurements. The electrolyte in the two reservoirs was electrically connected to an Axopatch 200B amplifier (Molecular Devices LLC, UK) by Ag/AgCl electrodes (silver wire chloridized in household bleach). Analog signals were digitalized by Digidata 1550B digitizer (Molecular Devices LLC, UK) and recorded on a computer with Clampex 10. 5 software (Molecular Devices LLC, UK). After flushing both chambers with DIW, the chambers were filled with 1 M KCl for current-voltage (I-V) measurement (voltages ranging from −120mV to 120mV). The diameter of nanopores was extracted from their conductance by using the simple model as described in Ref. ^1,10^. Unless stated differently, all measurements were performed under 500 kHz sampling, 100 kHz low-pass filter (four-pole internal Bessel filter), at room temperature of 21 °C.

### Lipid bilayer coating

In order to prevent the non-specific adsorption, surface passivation of the pore was implemented by using 1-palmitoyl-2-oleoyl-sn-glycero-3-phosphocholine (POPC, Avanti Polar lipids Inc., USA). Vials with POPC in chloroform were dried in a vacuum and subsequently stored at −20°C. Before usage, the lipids were re-suspended in 600KHM buffer (600mM KCl, 50mM HEPES, 5mM MgCl_2_, pH 7.5) to a concentration of 1mg/ml. The suspensions were then sonicated in a bath sonicator (Branson, model: 1510, USA) for >20min. Then, 50μL of lipid suspension was added to the ground-side reservoir of the nanopore, while applying an AC voltage with triangle waveform, 50 mV peak amplitude, and 1 Hz frequency. Lipid bilayer coating of the pore decreases the pore conductance. After stabilization for 10min, the entire flow cell was totally immersed in DIW and the chambers were flushed with DIW with further incubation for 20 min. Then, the chambers were flushed with DIW again before the flow cell was taken out of the bath and dried externally, filled with 600KHM buffer, and reconnected to the amplifier. Finally, the I-V curve was measured again, and compared with the I-V before coating, to extract the size of the nanopore before and after coating.

### DNA-origami spheres and protein samples

The DNA-origami sphere was designed based on ref.^1^ using cadnano2. The DNA-origami sphere was produced as previously described.^11^ Oligonucleotides and scaffold were purchased through tilibit nanosystems GmbH (Munich). Briefly, the folding mixture contained a 7560-nucleotide long single-stranded DNA scaffold at a final concentration of 50 nM, staple strands at a final concentration of 175 nM (3.5× fold excess), and folding buffer (5 mM TRIS, 1 mM EDTA, 5 mM NaCl and 20 mM MgCl_2_). The reaction mixtures were annealed in a T100 Thermal Cycler (Biorad) device using the following ramp: 65°C for 15 min, 60°C-40°C (1°C/hour). Afterwards, the reaction mixtures were incubated at room temperature. The reaction mixtures were purified by ultrafiltration, using Amicon Ultra 0.5 mL Ultracel filters (100k) and buffer containing 5 mM TRIS, 1 mM EDTA, 5 mM NaCl and 5 mM MgCl_2_. First, the filters with 500 μL of buffer were centrifuged for 5 min at room temperature (RT) at 10,000 g. After removing the filtrate, 50 μL of the sample was diluted with 450 μL of buffer and centrifuged again for 5 min at RT at 10,000 g. 450 μL of buffer was added to the filters, which were centrifuged for 5 min at RT at 10,000 g. This step was repeated 3 times. For retrieving the sample, the filter was removed, placed upside-down in a new tube, and centrifuged for 5 min at RT at 10,000 g. The cholesterol-modified oligos were added at 4× excess per handle (total: 24× excess) overnight. Afterwards, the sample was purified by ultrafiltration using the same protocol as described above. Models of the structure were computed with CanDo ^12^ and rendered with ChimeraX.^13^ In this study, docking was initiated by inserting 50μl of 20nM of DNA-origami sphere solution in 600KHM buffer into the cis chamber at positive voltage.

Avidin (from egg white) was purchased from Thermo Fisher Scientific (USA). ClpP was expressed and purified in-house as previously described.^14^ dCas9 was purchased from New England Biolabs Inc. (USA). Other proteins were purchased as Gel Filtration Calibration Kits from Cytiva LLC. (USA). All proteins were dispersed in 600KHM buffer at the indicated concentrations.

### Data processing

Data were analyzed using home-made code in MATLAB platform. For the detection of trapping events, the function *‘findchangepts’* was adopted to find the time points of the current level changes between the docking state and trapping state. Information about each trapping event, including the duration, blockage amplitude, and interval between the two adjacent events, could be extracted. In addition, an amplitude threshold for trapping event detection was placed at three times the standard deviation of the noise of the docking baseline. In order to estimate the uncertainties of kinetic rate constants (i.e. trapping time), bootstrap sampling was used: for a set of n data points, 10 subsets with a size of 60% of n were randomly picked with replacement, fitted by an exponential distribution separately. Then, the means and standard deviations of the extracted trapping time across these subsets are calculated.

### COMSOL simulations

The numerical simulations of the NEOtrap system were implemented on COMSOL Multiphysics 5.4 with a two-dimensional axial symmetrical domain. The simulations included the fluid, the membrane, and the DNA-origami sphere, whose relative permittivity was set to 80, 7.5,^15^ and 8.3,^16^ as deduced for water, SiNx, and DNA, respectively. The ion distribution and movement in an electrolyte were governed by the Nernst-Planck equation, the electric potential distribution was described by the Poisson equation, and the fluid flow was determined by the Navier-Stokes equations. The *Transport of Diluted Species* module (Nernst-Planck equation), the *Electrostatics* module (Poisson equation), and the *Laminar Flow* module (Navier-Stokes equations) were incorporated and fully coupled in the simulation. The electrolyte was 600 mM KCl with the mobilities of K^+^ and Cl^-^ were 7.0 × 10^-8^ and 7.2 × 10^-8^ m^2^ V^-1^ s^-1^, respectively.^17^ The respective diffusion coefficient was then determined through the Einstein relation. It is worth noting that in order to reduce the demand for computation resources and to reach a converged model, a two-dimensional axial symmetry was adopted instead of a full three-dimensional simulation. Strictly speaking, however, neither the vertical nor the horizontal configurations of the origami sphere possess axial-symmetry. Yet, the two-dimensional simulations do capture the major differences between the vertical and horizontal configurations.

## ASSOCIATED CONTENT

### Supporting Information

The Supporting Information is available free of charge at https://pubs.acs.org/doi/XXX.

This SI contains: Docking of the bare or cholesterol-functionalized origami sphere onto pores; Estimation of the viscous force on a trapped protein; The trapping energy well; Current traces of docking of cholesterol-functionalized DNA origami spheres under a negative voltage ramp; Current traces of the undocking of bare DNA-origami spheres; Current traces of the open-pore with lipid bilayer coating; Current trace showing docking and undocking of a cholesterol-functionalized DNA-origami sphere; Current traces showing the controlled docking of DNA-origami spheres in a vertical or horizontal orientation; Trapping data of ovalbumin; Trapping time of avidin proteins at different voltages by bare and cholesterol-functionalized DNA-origami spheres; Trapping Ribonuclease A by using a cholesterol-functionalized DNA-origami sphere or bare DNA-origami sphere; Current traces showing the trapping of different proteins by a bare origami sphere and a vertically locked cholesterol-functionalized origami sphere. (PDF)

## Acknowledgements

We thank Hendrik Dietz and Pierre Stömmer for discussions and an early cholesterol functionalization design, and Frans Tichelaar for TEM drilling. Avidin was a kind gift of Mark Howarth. ClpP plasmids were a kind gift of Chirlmin Joo. The work was funded by NWO-I680 (SMPS) and supported by the NWO/OCW Gravitation program NanoFront and the ERC Advanced Grant 883684.

## Notes

The authors declare no competing financial interest.

## Data availability

Data are available at https://doi.org/10.5281/zenodo.7047595.

---

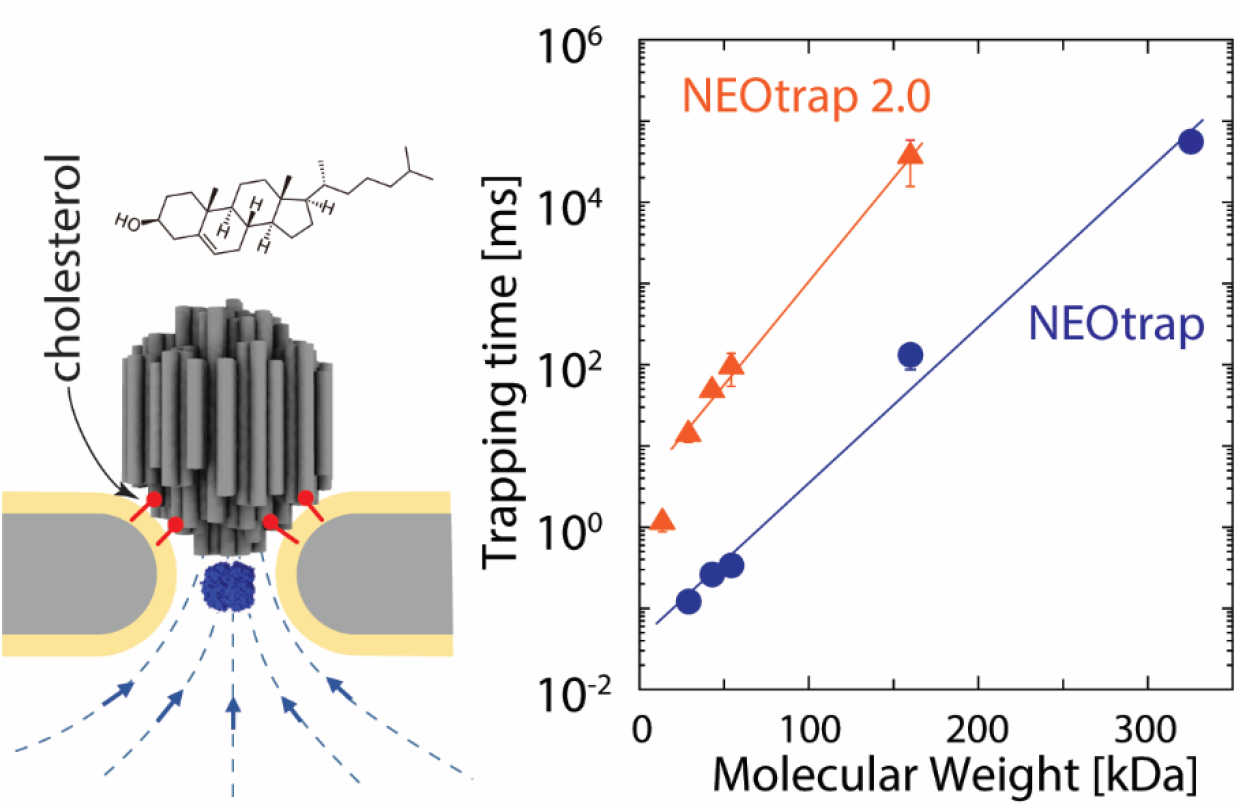

