## Supporting Information for "Orientation-locked DNA origami for stable trapping of small proteins in the NEOtrap"

**Table of Contents**

Note 1. Docking of the bare or cholesterol-functionalized origami sphere onto pores

Note 2: Estimation of the viscous force on a trapped protein

Note 3. The trapping energy well

Figure S1. Current traces of docking of cholesterol-functionalized DNA origami spheres under a negative voltage ramp

Figure S2. Current traces of the undocking of bare DNA-origami spheres

Figure S3 Current traces of the open-pore with lipid bilayer coating

Figure S4. Current trace showing docking and undocking of a cholesterol-functionalized DNA-origami sphere

Figure S5. Current traces showing the controlled docking of DNA-origami spheres in a vertical or horizontal orientation

Figure S6. Trapping data of ovalbumin.

Figure S7. Trapping time of avidin proteins at different voltages by bare and cholesterol-functionalized DNA-origami spheres

Figure S8. Trapping Ribonuclease A by using a cholesterol-functionalized DNA-origami sphere or bare DNA-origami sphere

Figure S9. Current traces showing the trapping of different proteins by a bare origami sphere and a vertically locked cholesterol-functionalized origami sphere

**Supporting Note 1: Docking of the bare or cholesterol-functionalized origami sphere onto pores.**

In order to measure the strength of the interaction between the cholesterol anchors and the lipid bilayer, we linearly decreased the voltage and recorded the corresponding current traces. As shown in Fig. S1(a), docking of a single origami sphere happened in a holding period with a constant 100 mV bias. After docking, the voltage was ramped down from 100 mV to -500 mV. Beyond a certain negative voltage, the noise suddenly increased, indicating that the lipid bilayer became unstable, and the sphere undocked under the high electric field. No distinct changes were observed until this voltage (in contrast to the data for bare pores, cf. Fig. S3). The voltages causing the instability of the lipid bilayer were regarded as the release voltage of the cholesterol-functionalized origami spheres.

Compared to the cholesterol-functionalized origami sphere, the bare origami spheres displayed different current traces during a negative voltage ramp. As shown in Fig. S2, a sudden current increase was observed at about positive +50 mV, which was caused by the undocking of the sphere whereupon the current returned back to the open pore level. Thus, for a bare origami sphere, even a small positive voltage was insufficient to stably hold the sphere on the pore, which we attribute to thermal fluctuations.

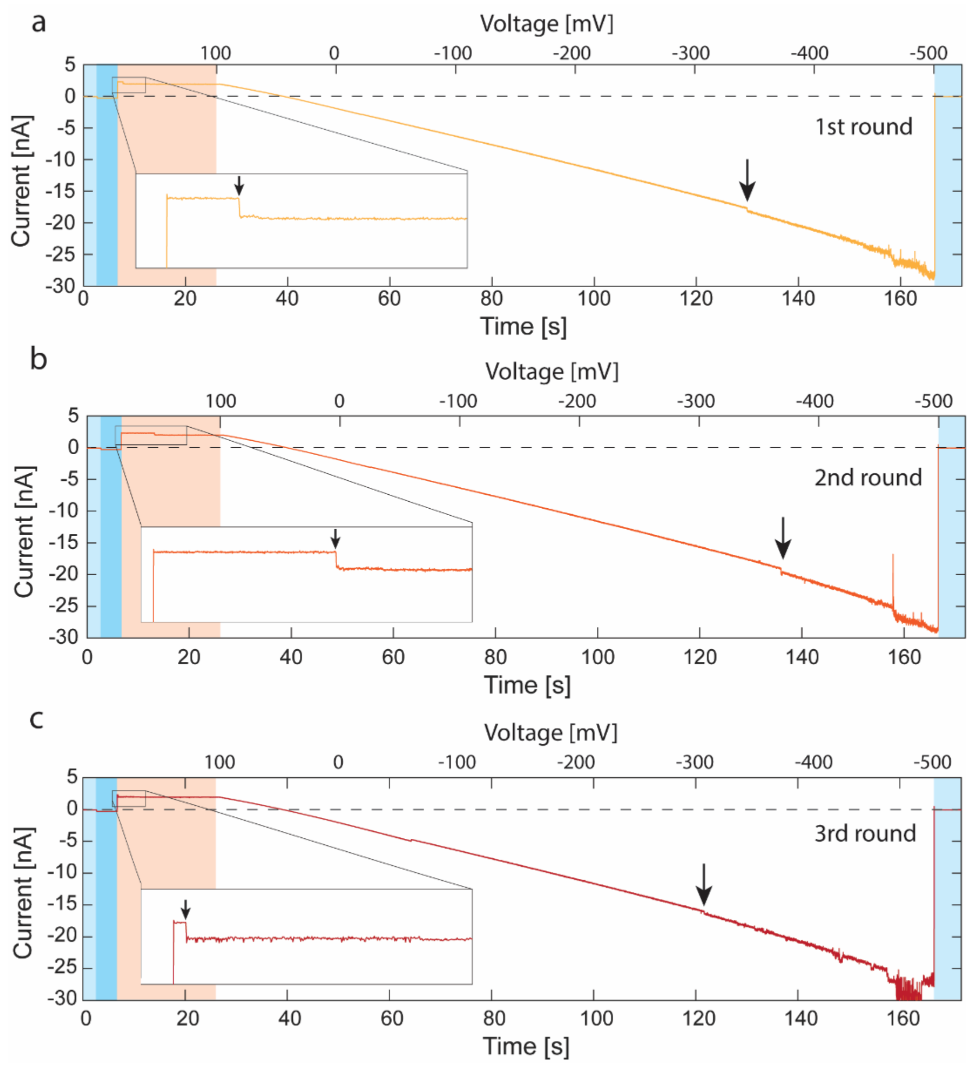


**Figure S1. Current traces of the docking and undocking of cholesterol-functionalized DNA-origami spheres under a negative voltage ramp.** (a-c) Traces of the 1st, 2nd, and 3rd rounds of the voltage cycle. In the light blue region, the bias voltage was zero. Then, -20 mV was applied to repel any possible docked sphere, marked as the dark blue color. Afterwards, +100 mV bias was added for 15s to attract origami spheres and dock one sphere onto the nanopore, for a period marked in the orange color. In this period, a current drop was observed, which is caused by the docking of an origami sphere due to the steric blockage of the pore, as indicated by the arrow in the zoomed-in view of the inset. Next, the bias voltage was ramped down linearly from 100 mV to -500 mV. A gradual change of current was observed correspondingly. At a certain negative voltage as marked by the arrow, the current noise suddenly increased. We attribute this to a sudden instability of the lipid bilayer induced by the high electric field, and the point where the cholesterol anchors of the origami sphere are likely pulled out from the lipid bilayer to release the sphere which is electrophoretically pulled away from the pore. In the next round, the lipid bilayer resealed and a new docking by another origami sphere could be studied. A similar pattern appeared every round of the repeated cycles, as shown in (a) to (c).

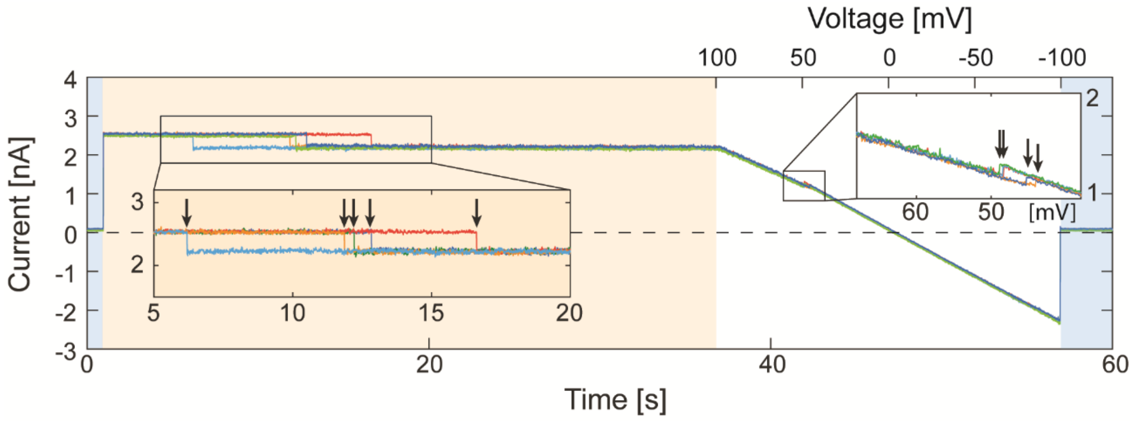


**Figure S2. Current traces of the undocking of bare DNA-origami sphere.** Voltage cycles were repeated five times and the corresponding current traces are shown with different colors. In the light blue region, the bias voltage was zero. Then, +100 mV bias was added for 36s (orange color) to attract an origami sphere and dock it on the nanopore, leading to the observed current drop, as indicated by the arrows in the zoomed-in view of the left inset. Afterwards, the bias voltage was ramped down gradually from 100 mV to -100 mV, and a gradual decrease in the current was observed. At a certain voltage of about +50 mV (marked by the arrows in the zoomed-in view of the right inset, the horizontal axis is voltage) the current suddenly increased, indicating the undocking of the sphere. A similar pattern was found in every round of the voltage cycles.

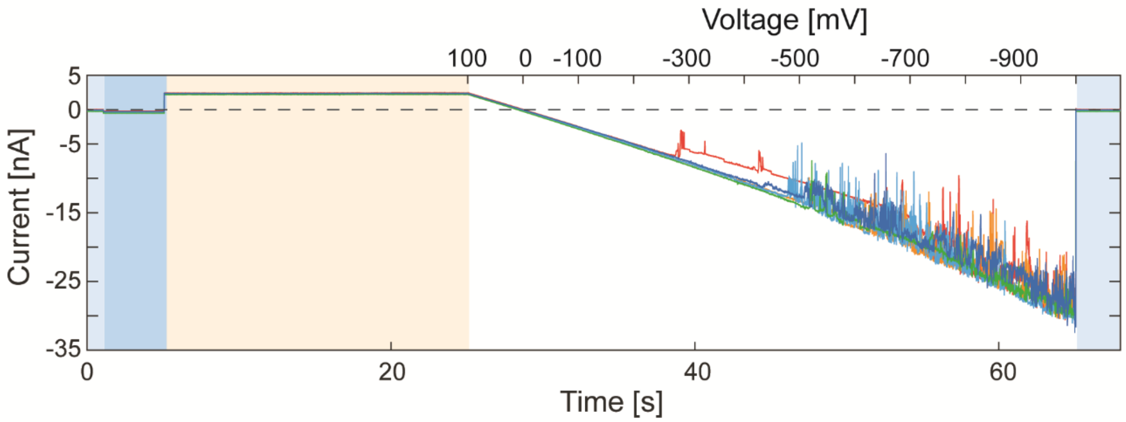


**Figure S3 Current traces of the open-pore with lipid bilayer coating under a negative voltage ramp.** Voltage cycles were repeated five times and the corresponding current traces are shown in different colors. The voltage cycle was similar to that in Figure S1. The only quantitative difference was that the voltage decreased from 100 mV to -1000 mV. Similar to the traces in Figure S1, the noise suddenly increased at a certain negative voltage, indicating an unstable lipid bilayer. This suggests that the origami sphere locked by the cholesterol molecules is only able to undock if the lipid bilayer becomes unstable.

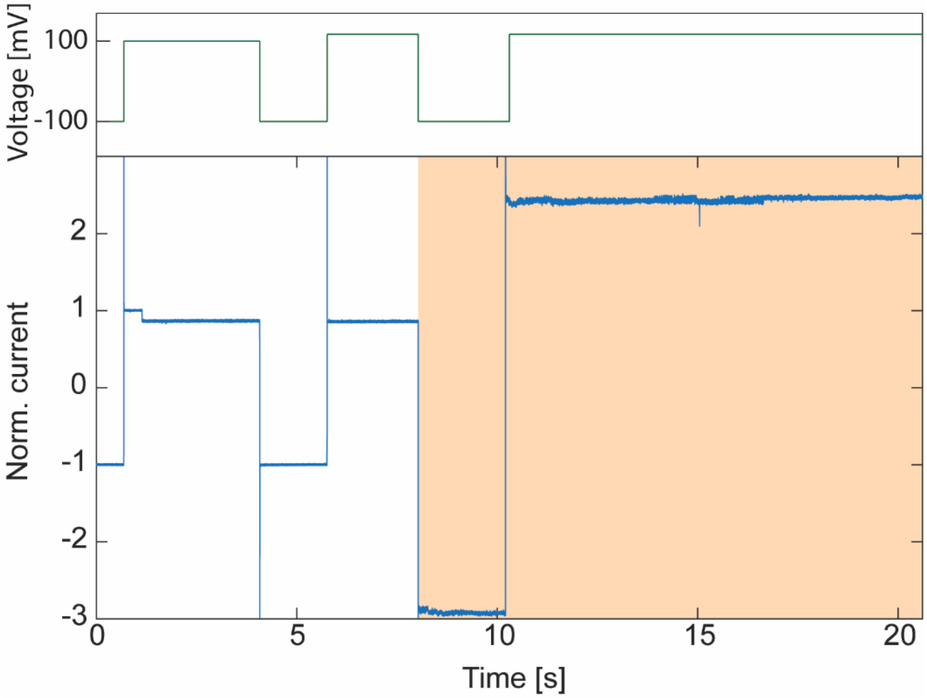


**Figure S4. Current trace showing docking and undocking of a cholesterol-functionalized DNA-origami sphere.** The bottom panel shows the current versus time. The corresponding profile of the applied voltage is shown in the upper panel. When a 100 mV bias was applied, a current decrease step was found, indicating the docking of an origami sphere. Since the cholesterols anchored the sphere onto the nanopore, the current was kept at the docking level upon switching the voltage to ±100 mV. Finally (orange region), application of -100 mV bias removed the sphere and peeled off the lipid bilayer, yielding a very high current that indicates a bare pore without a lipid bilayer.

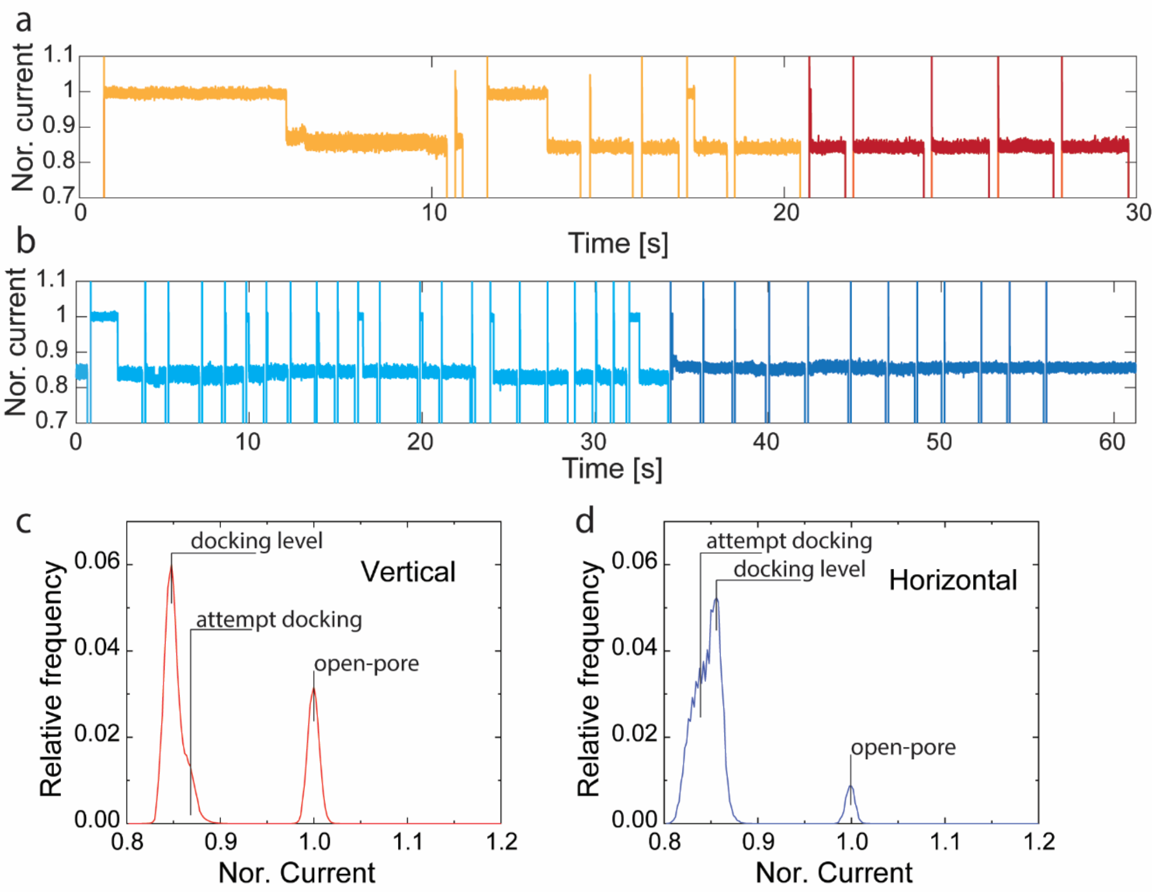


**Figure S5. Current traces showing the controlled docking of DNA-origami spheres in a vertical or horizontal orientation.** (a) Current trace of vertical sphere docking. Here, cholesterol molecules were only functionalized at the bottom of the origami sphere. As shown in yellow color, a current drop was observed after every voltage inversion in a series of alternating voltages, indicating the sphere was not well locked by the cholesterol. However, in the voltage cycle at 21 s (red color), the sphere was docked 'irreversibly' (i.e. presumably stably locked by cholesterol), in such a way that the negative voltage did not remove the sphere. Instead, the current stayed at the docking level after following voltage inversions. (b) Similar current traces for horizontal sphere docking. The sphere was locked by the cholesterols in the voltage cycle at 34 s whereupon afterwards, the current stayed at the docking level shown in the dark blue color. (c) Current distribution of the vertical docking, as deduced from the trace in panel a. (d) Same as c but for horizontal, based on trace in panel b. For the vertical configuration, the current level of stable docking locked by cholesterols was lower than the level of attempt temporary docking. The reversed phenomenon was observed for the horizontal configuration, i.e., the locked docking level was higher than the attempt docking level. We attribute this to the non-isotropic sphere geometry which allows the vertically oriented sphere to reach deeper into the pore – and thus block more through-pore current – than the horizontally oriented sphere.

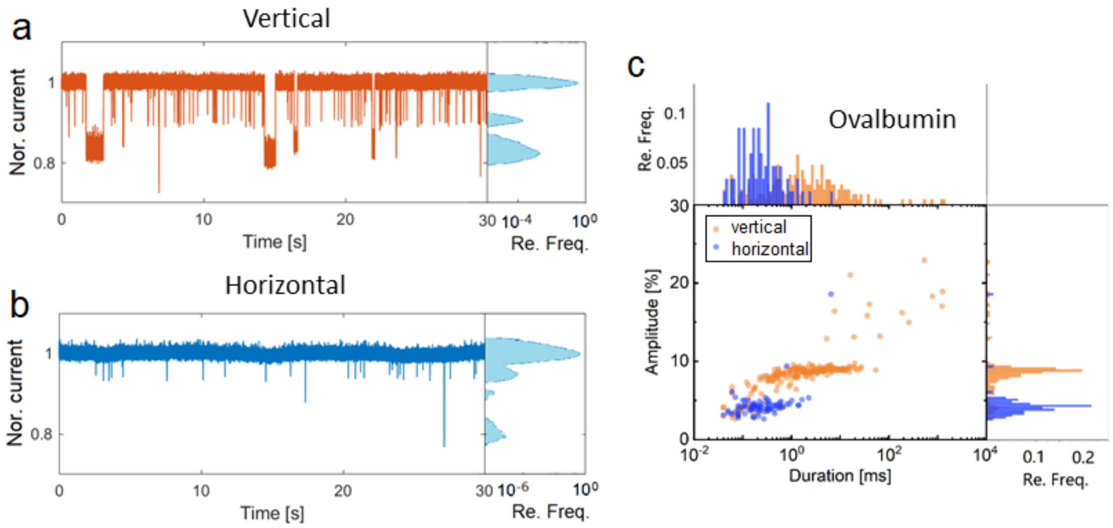


**Figure S6. Trapping date of ovalbumin.** Current traces show the trapping of ovalbumin by vertically (a) and horizontally (b) docked origami spheres at 100 mV bias. The histograms of the corresponding current traces are shown on the right. (c) Scatter plots compare the trapping time and relative blockage amplitude of the trapping events from panels a and b.

**Supporting Note 2: Estimation of the viscous force on a trapped protein**

According to Stokes' law, a viscous sheer force acting on a sphere can be calculated by

$$F_v = 6\pi\eta v_{eof} r$$

where $\eta$ is the viscosity of water, $v_{\mathrm{oef}}$ is the water velocity, and $r$ is the radius of the sphere. Approximating the shape of proteins by a sphere, the viscous force to hold a trapped protein in the pore can thus be estimated.

From the simulation, the maximum velocity of the EOF in the pore is around 0.1 m/s for the vertically locked origami sphere (Figure 3b in the main text). The radius of an avidin molecule is ~3 nm. Thus, the approximate viscous force to hold the trapped avidin is in the single-digit pico-newton range:

$$F_v = 6\pi \times 10^{-3}[Pa \cdot s] \times 0.1[m/s] \times 3 \times 10^{-9}[m] = 5.6[pN]$$

**Supporting Note 3: The trapping energy well**

The escape from the trapping potential well can be described as an energy barrier-crossing process. The escape rate $k_{\mathrm{off}}$ (*i.e.*, the reciprocal of the trapping time $\tau_{\mathrm{trap}}$) then follows an Arrhenius relationship with the energy barrier height,

$$k_{off} = \frac{1}{\tau_{trap}} = k_0 \exp(-\frac{E_b}{k_B T}) \quad ,$$

where, $k_0$ is a rate constant, $E_{\mathrm{b}}$ is the trapping energy barrier height, $k_B$ is the Boltzmann constant, and $T$ is the absolute temperature. Thus, a prolonging of the trapping time indicates the increase of the energy barrier of escape,

$$A = \frac{\tau_{trap,2}}{\tau_{trap,1}} = \exp(\frac{E_{b,2} - E_{b,1}}{k_B T}) = \exp(\frac{\Delta E}{k_B T}) \quad ,$$

where $A$ is the ratio of prolonged trapping, and $\Delta E$ the corresponding increase of the barrier height. An increase of the trapping time by a factor 100 thus means an increase of 4.6 $k_{\mathrm{B}}T$ in barrier height.

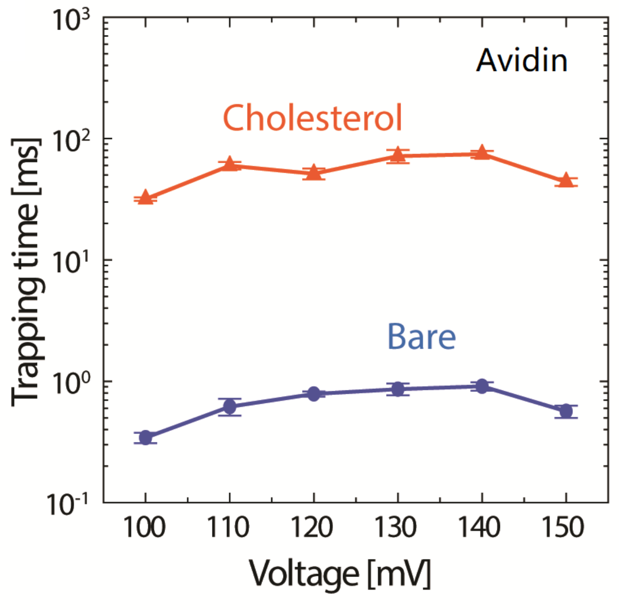


**Figure S7. Trapping time of avidin proteins at different voltages by bare and cholesterol-functionalized DNA-origami spheres.**

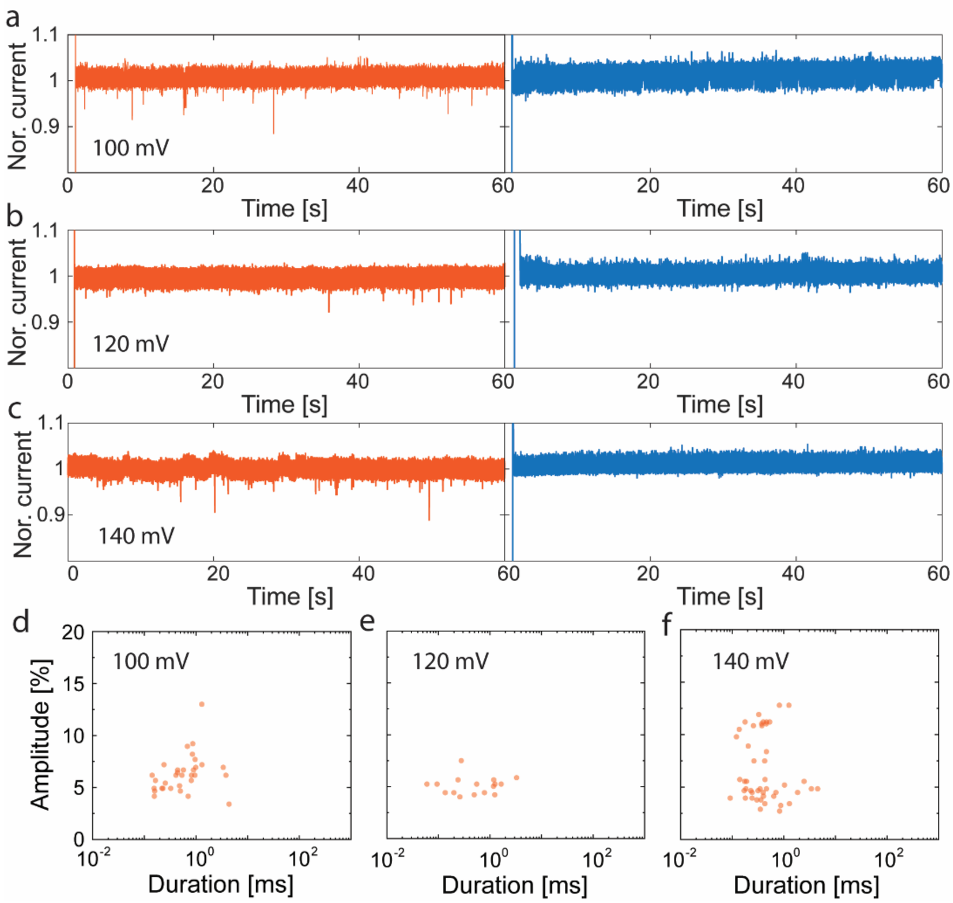


**Figure S8. Trapping of Ribonuclease A (13.7 kDa) using a cholesterol-functionalized or bare DNA-origami sphere.** (a-c) Typical current traces of trapping by a cholesterol-functionalized origami sphere (left panel) or a bare origami sphere (right panel) at 100 mV, 120 mV, and 140 mV, respectively. (d-f) Scatter plots showing the trapping time and relative blockage amplitude of the trapping events of Ribonuclease A for cholesterol-functionalized origami spheres at different voltages from (a) to (c). No trapping events were observed with the bare origami sphere.

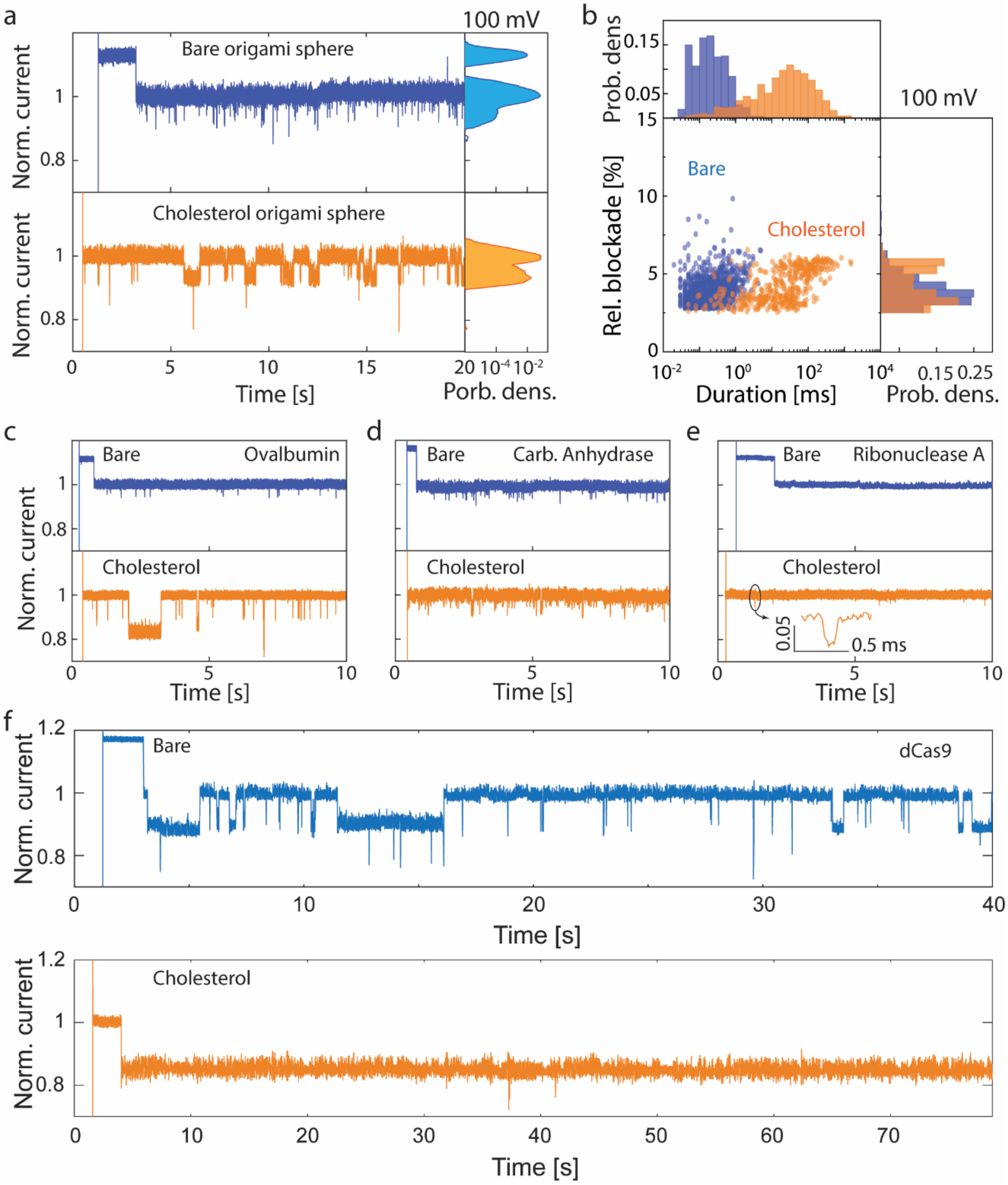


**Figure S9. Current traces showing the trapping of different proteins by a bare origami sphere and a vertically locked cholesterol-functionalized origami sphere at 100 mV bias.** (a) Trapping of avidin at 100 mV. Histograms of the corresponding current traces are shown at the right-hand side. (b) Scatter plot that compares the trapping time and relative blockage amplitude from data of panel a. (c-f) Current traces of trapping ovalbumin (c), carbonic anhydrase (d), ribonuclease A (e), and dCas9 (f) for both bare origami spheres (blue) and cholesterol-functionalized origami spheres (orange) at 100 mV.
